# Descending GABAergic Neurons of the RVM That Mediate Widespread Bilateral Antinociception

**DOI:** 10.1101/2023.04.29.538824

**Authors:** Robert P. Ganley, Marília Magalhaes de Sousa, Matteo Ranucci, Kira Werder, Tugce Öztürk, Hendrik Wildner, Hanns Ulrich Zeilhofer

## Abstract

Neurons projecting from the rostral ventromedial medulla (RVM) to the spinal dorsal horn are critical elements of endogenous pain control systems. Here, we describe a GABA/glycinergic pathway that predominantly innervates the superficial dorsal horn. Anatomical and optogenetic tracing of these neurons from a single unilateral site of the lumbar spinal cord indicated that these neurons give rise to a dense bilateral innervation of the spinal cord along its entire rostrocaudal axis. Chemogenetic activation of these neurons caused a bilateral and widespread reduction in heat, cold, and mechanical sensitivity, while their silencing with tetanus toxin induced allodynia and spontaneous pain-like aversive behaviors. Consistent with a continuous role in the prevention of spontaneous pain, many descending RVM GABAergic neurons were found to be tonically active. This pathway may therefore be relevant for widespread conditioned analgesia, while its dysfunction may underlie chronic widespread pain syndromes.

## Introduction

The rostral ventromedial medulla (RVM), which includes the nucleus raphe magnus (NRM), the nucleus paragigantocellularis pars alpha (Pa) and the lateral paragigantocellularis (LPGi), is a region within the ventral hindbrain critically involved in endogenous pain modulation. The ability of this region to strongly suppress nociception has been established through stimulation-produced analgesia experiments, in which stimulation of certain brain regions, such as the periaqueductal grey (PAG) strongly suppressed nociception ^1^.

Descending projections from the RVM are especially important for context-dependent regulation of pain sensitivity. This system requires intact descending fibers within the dorsolateral funiculus since lesioning these tracts removes most of the antinociceptive effects of stimulation-produced analgesia ^2^. Although many projection neurons and neurotransmitters of these descending system have been identified, their precise anatomical organisation and physiological functions are only incompletely understood.

Much of our understanding of the functional role of RVM neurons is based on pharmacological and physiological characterization of neurons with single unit recordings. Recorded neurons are classified as either ON, OFF, or Neutral cells, depending on changes in their activity in response to acute nociceptive stimulation ^3^, and their activities are thought to initiate the withdrawal from damaging stimuli. The activity patterns of ON and OFF neurons are modulated by input from the lateral parabrachial area (LPb)^4^, and are known to change during and following inflammatory hyperalgesia ^5, 6^. Besides their responses to noxious stimuli and pharmacological agents little is known about the identity of these pro-and antinociceptive types, such as their neurotransmitter content and their influence on pain-related behaviors in awake animals. Further, it is unclear whether these neurons are local interneurons or project their axons to the spinal cord to directly suppress the transmission of nociceptive information to higher brain centers.

Descending pain control involves multiple transmitter systems and many separate neuron populations ^7–9^. The RVM provides a particularly strong innervation of the spinal cord and controls pain sensitivity in a bidirectional manner ^8, 10^. Many of the descending projection neurons in the RVM are accessible with viral tools, which provides an opportunity to study them with greater precision ^11–14^. This is particularly useful for selectively targeting projection neurons containing neurotransmitters that are present in many regions the nervous system, such as GABA, glycine, and glutamate, as it is challenging to restrict gene expression only to the projection neurons within a given region. Selective gene expression in a neuronal population is often a prerequisite for functional studies, since these require the restricted expression of receptors, ion channels, and toxins, which is commonly achieved in neuroscience with viral tools and transgenic animals^15^.

We aimed to study the functional roles played by inhibitory neurons that project directly to the spinal cord from the RVM. Together with vGAT^cre^ mice we utilized AAV2retro vectors containing Cre-dependent constructs to dissect this pathway. We found that activation of this pathway produces widespread antinociception through wide-ranging axonal projections throughout the rostrocaudal extent of the spinal cord. In contrast, when these neurons are silenced mechanical sensitivity to dynamic and punctate stimuli is strongly increased. These results provide an extensive anatomical and functional dissection of this pathways under physiological conditions, and demonstrate the potential of these projections for pain suppression at the spinal cord.

## Results

### Specific labelling of descending inhibitory RVM projections using AAV2retro vectors in vGAT^cre^ mice

To study the descending pathways from the RVM we used AAV2retro vectors to transduce RVM projection neurons via their axon terminals in the dorsal horn^12^. We injected AAV2retro.eGFP into the left lumbar dorsal horn of 9–10-week-old male wild-type mice, and prepared the hindbrain tissues for multiplex fluorescent *in situ* hybridization (fISH) for vGAT and vGluT2 mRNA (Figure 1A). We found that many eGFP-labelled cells contained vGAT (19.4 ± 4.1%) or vGluT2 (55.1 ± 7.0%) mRNA. Therefore, to restrict expression to the inhibitory subset we performed experiments with vGAT^cre^ mouse driver lines to address the function of this population. As a first step, we confirmed eutrophic cre expression in vGAT^cre^ mouse lines in descending RVM neurons. We injected AAV2retro vectors that contained a Cre-dependent eGFP sequence (AAV2retro.flex.eGFP) into the lumbar dorsal horn and examined the hindbrains with multiplex fISH (Figure 1B). The majority of eGFP+ hindbrain neurons in vGAT^cre^ animals contained vGAT mRNA (76.0 ± 16.6%) with few containing detectable vGluT2 (7.3 ± 4.7%). This restricted expression of eGFP in vGAT^cre^ mouse lines confirms that inhibitory descending neurons can be selectively labelled from the spinal cord using AAV2retro vectors containing Cre-dependent constructs.

**Figure 1:**
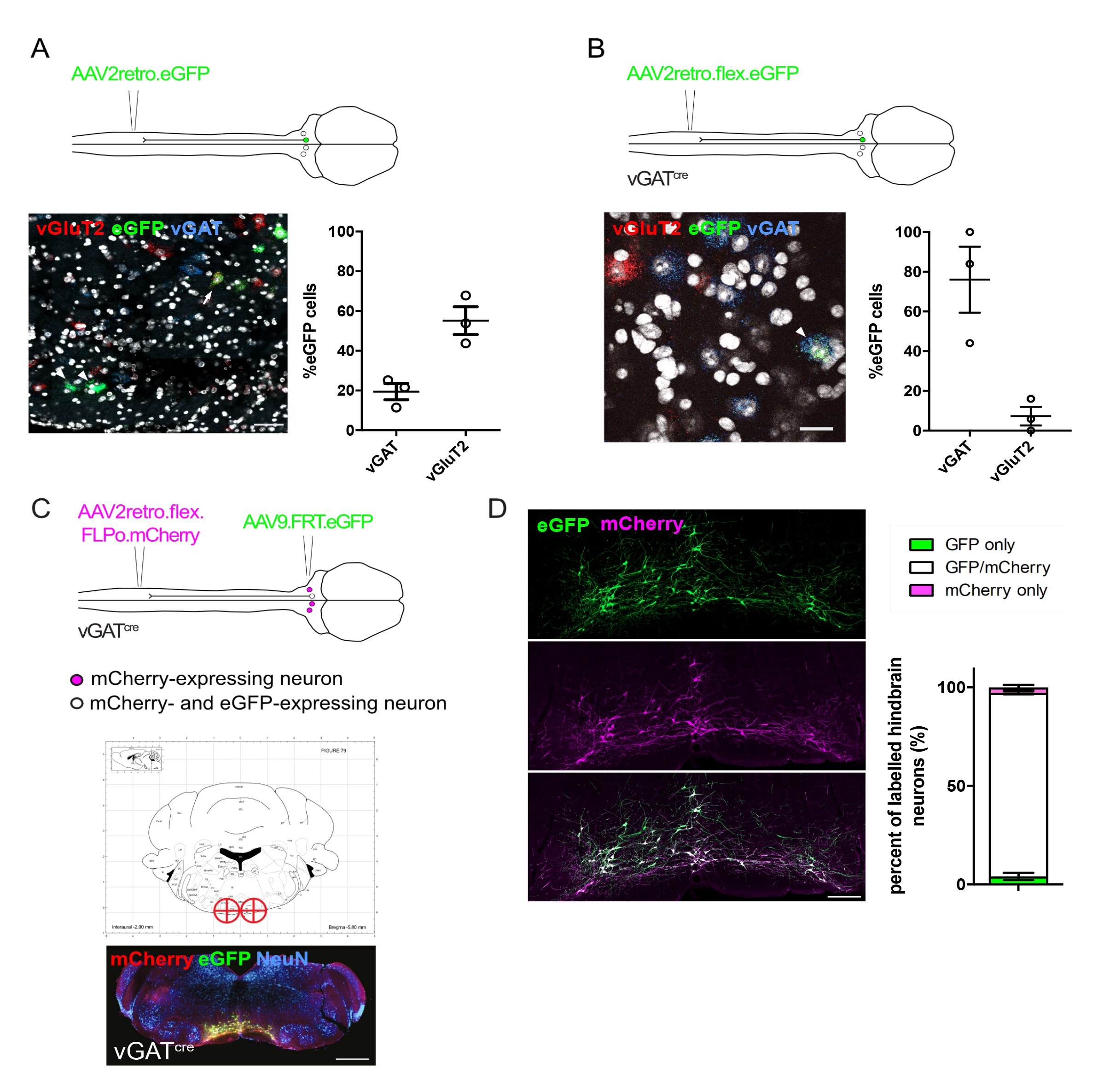
RVM projection neurons labelled with AAV2retro from the spinal cord and intersectional strategy for targeting inhibitory descending projection neurons. A. Labelling descending projection neurons with AAV2retro. Fluorescent *In situ* hybridization of an RVM section showing eGFP+ neurons that colocalize with vGAT or vGluT2 mRNA (arrowheads or arrows respectively)(scale bar = 20 μm). Quantification of eGFP+ neurons that only colocalized with vGAT or vGluT2 mRNA (n = 3 animals). B. Labelling descending vGAT^cre^ RVM neurons. Fluorescent *In situ* hybridization of RVM sections taken from vGAT^cre^ animals that received an intraspinal injection of AAV2retro.flex.eGFP (scale bar = 10 μm). Quantification of eGFP+ cells in the RVM containing vGAT or vGluT2 mRNA (n = 3 animals). C. Intersectional strategy for specifically targeting spinally-projecting inhibitory neurons of the RVM. Brain injection coordinates for the RVM injections and representative injection sites from labelling experiments (scale bar = 200 μm in both images). D. Higher magnification image of mCherry+ and eGFP+ labelled neurons in the ventral hindbrain (scale bars = 100 μm). Quantification of mCherry-and eGFP-expressing neurons in the RVM of vGAT^cre^ animals (n = 3 animals)

Since vGAT^cre^ is expressed in the spinal cord as well as the RVM, we devised an intersectional approach to specifically study GABAergic descending projection neurons without also targeting local spinal or RVM neurons ^16, 17^. To this end, the lumbar spinal cords of these animals were injected with AAV2retro vectors containing an optimized Cre-dependent flippase (Flpo), and one week later, the RVM of these animals was injected with a Flpo-dependent reporter (AAV9.dFRT.eGFP) (Figure 1C).To assess the specificity of this approach, we also included an mCherry construct in the AAV2retro (AAV2retro.flex.FLPo.mCherry) to directly label the projections, which in turn allowed us to determine the accuracy and completeness of the hindbrain injections with AAV9.dFRT.eGFP.

We optimized the brain injection coordinates to label most vGAT^cre^ projection neurons of the RVM, based on their location and distribution in the brain stem (Figure 1C). The vGAT^cre^ projection neurons were located both in medial and lateral regions of the RVM, which required bilateral injections to label efficiently (−5.8, ±0.5, 5.9 from Bregma). The majority of labelled cells contained both mCherry and eGFP (93.1 ± 0.8%). Few neurons contained mCherry or eGFP alone (2.8 ± 1.3% and 4.1 ± 1.8% respectively), highlighting both the specificity and sensitivity of this approach (Figure 1D). Taken together, the viral tools and the labelling strategy provided reliable means of accessing these RVM descending neurons for functional and anatomical studies.

### Global wide-ranging projections of descending inhibitory and excitatory neurons

The spinal dorsal horn has a structured laminar organization, which reflects the segregated innervation by sensory fibers of different sensory modalities ^18, 19^. Hence, the spinal termination area of descending projections will likely be important for their function. We therefore characterized the axonal projection pattern of the descending neurons (Figure 2A). Within the spinal cord, the inhibitory descending neurons terminated in particular laminar regions (Figure 2B). Inhibitory terminals were densely located in the superficial laminae with only sparse innervation of the rest of the spinal cord. Bundles of axons were also present ventral to the superficial spinal laminae within laminae IIi – V.

**Figure 2:**
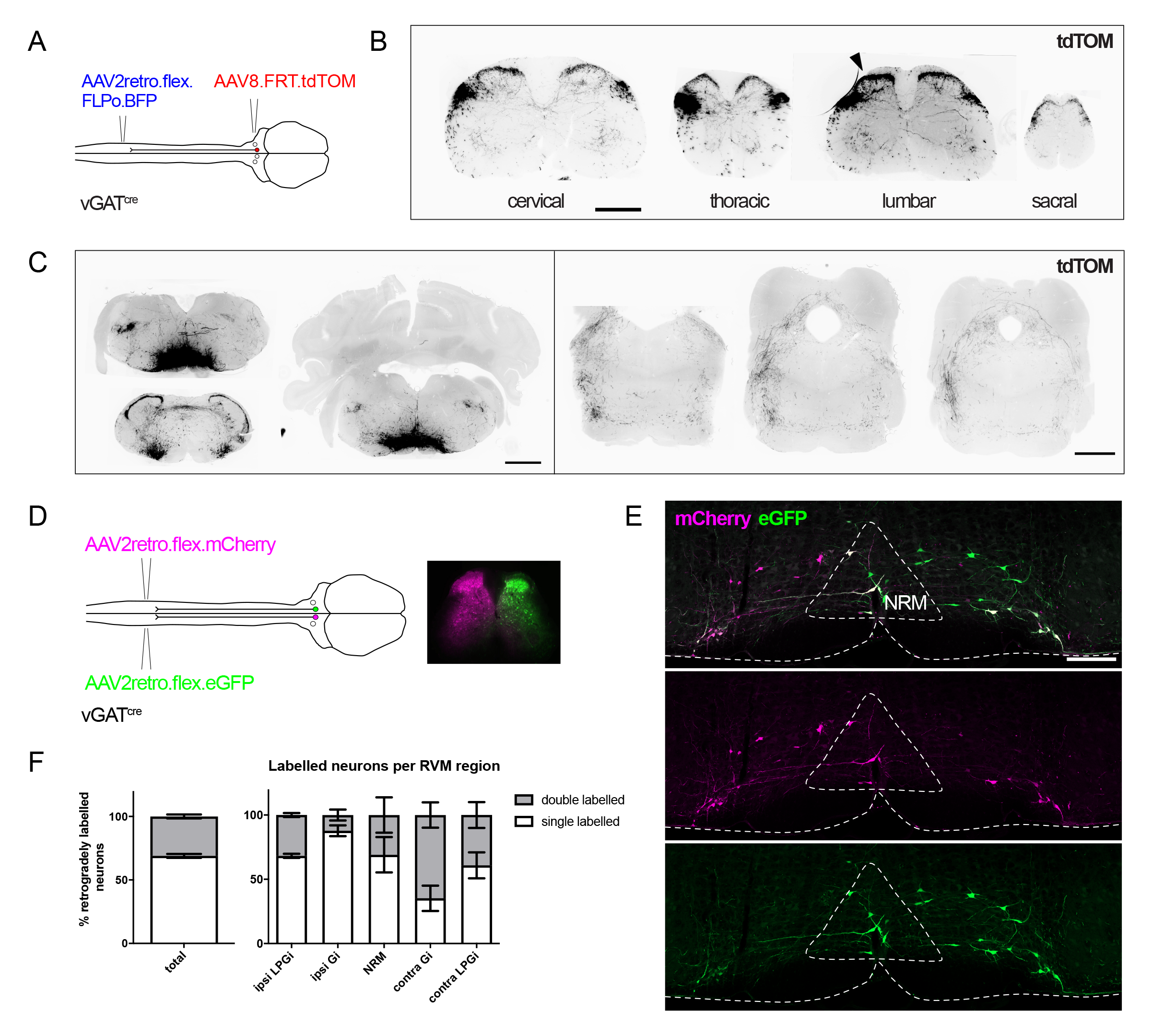
Anatomical tracing of descending inhibitory RVM neurons: A. Injection scheme for labelling descending RVM projection neurons with tdTOM. B. Spinal cord sections illustrating the axon terminal pattern of descending vGAT^cre^ projection neurons of the RVM. Arrowhead indicates the injection site of the AAV2retro.flex.FLPo.BFP in the left lumbar spinal cord but axon collaterals are present in multiple spinal segments bilaterally (scale bar = 200 μm). C. injection site and rostral branches arising from vGAT^cre^ and RVM descending neurons. Axons from vGAT^cre^ neurons are found in the spinal trigeminal nucleus, the lateral parabrachial area (LPb) and periaqueductal grey matter (PAG)(scale bars = 500 μm). D. Injection scheme for retrogradely labelling RVM projection neurons from the left and right spinal cord. Representative injection sites from a vGAT^cre^ animal that received bilateral spinal cord injections showing the spread of each AAV is largely restricted to the injected side (scale bar = 100 μm). E. Neurons labelled in the hindbrain from the injections illustrated in D (scale bar = 500 μm). F. Quantification of vGAT^cre^ hindbrain neurons labelled with either one (unilateral) or two fluorophores (bilateral). The proportion of vGAT^cre^ neurons in different regions of the RVM are shown, illustrating that a larger proportion of unilateral neurons project to the ipsilateral spinal cord. N = 6 animals

Strikingly, labelled axon terminals were not only found at the spinal cord injection site (lumbar enlargement left side), but also on the contralateral side and throughout all spinal segments and including the medullary dorsal horn (Figure 2B and C). These experiments also gave us the opportunity to investigate the potential presence of rostrally projecting collaterals. We found that inhibitory neurons retrogradely labeled from the lumbar dorsal horn gave rise to axon collaterals that projected rostrally from the cell somata, and innervated multiple brain regions (Figure 2C). Axon collaterals from the vGAT^cre^ neurons innervated multiple sensory information processing CNS areas including the trigeminal nucleus, the nucleus of the solitary tract (NTS), the lateral parabrachial area (LPb), and the periaqueductal grey matter (PAG). Notably, many of these areas are part of the ascending pain pathways in rodents, representing termination zones of ascending spinal projections ^20, 21^. To visualize this elaborate projection pattern in 3D space, we repeated the labelling experiments in vGAT^cre^ animals and prepared nervous tissues for CLARITY-based tissue clearing, and imaged transparent brain tissue using light sheet microscopy ^22^. This labelling in transparent tissue clearly shows axons that emanate from the cell body in both rostral and caudal directions and on both ipsilateral and contralateral sides of the nervous system (video 1).

To further quantify the extent to which descending RVM neurons projected to both sides, we injected two AAV2retro vectors containing either Cre-dependent mCherry or eGFP into the left or right lumbar dorsal horns (Figure 2D). Spread of the AAVs remained restricted to the injected side (Figure 2D). We found however that many retrogradely labelled descending vGAT^cre^ neurons were labelled from both sides, i.e. coexpressed mCherry and eGFP (69 ± 1.5%), indicating that many of the descending RVM neurons projected to both sides of the spinal cord (Figure 2D and E). Of those cells that were labelled with only one fluorophore, most were generally found on the ipsilateral side of the RVM (60.2 ± 4.6%) (Figure 2F).

### Chemogenetic activation of descending inhibitory RVM projection neurons reduces cutaneous sensitivity

Our tracing experiments in vGAT^cre^ mice revealed dense innervation of the superficial dorsal horn laminae (Figure 2B). Since this area is the first node in the ascending pain pathway it is likely that this inhibitory innervation controls pain and pain-related behaviors ^19^. To assess the impact of this pathway on nociception, we performed chemogenetic activation experiments, introducing hM3Dq to the descending inhibitory neurons using our intersectional strategy (Figure 3A). The expression pattern of hM3Dq-mCherry was similar to our previous labelling experiments, with mCherry-labelled axon terminals present on both the injected and non-injected sides of the spinal cord (Figure 3B). Upregulation of the activity-dependent marker c-Fos verified the efficacy of CNO (vehicle: 41.2% hM3Dq neurons vs CNO: 67.18% p < 0.01 unpaired t-test) (Figure 3C and D). In agreement with findings of others^23, 24^, the activation of this descending pathway reduced the sensitivity of the ipsilateral paw to heat and cold stimuli, with withdrawal latencies increasing from 9.3 ± 0.9 to 14.5 ± 1.4 s in the Hargreaves assay and from 10 ± 0.9 to 14 ± 1.1 s in the cold plantar assay (Hargreaves vehicle vs CNO, p = 0.0011 paired t-test; cold plantar assay vehicle vs CNO p = 0.0089 paired t-test, Figure 3E)^23, 24^. In addition, activation of the vGAT^cre^ descending neurons reduced sensitivity to von Frey mechanical stimulation (from 4.34 ± 0.3 to 5.3 ± 0.4 g), indicating that this pathway is capable of suppressing sensitivity to multiple sensory modalities (von Frey: vehicle vs CNO p = 0.0045), Figure 3E)^23^. These changes in sensitivity were not accompanied by any detectable change in sensorimotor coordination (Rotarod: vehicle vs CNO p = 0.699 paired t -test).

**Figure 3:**
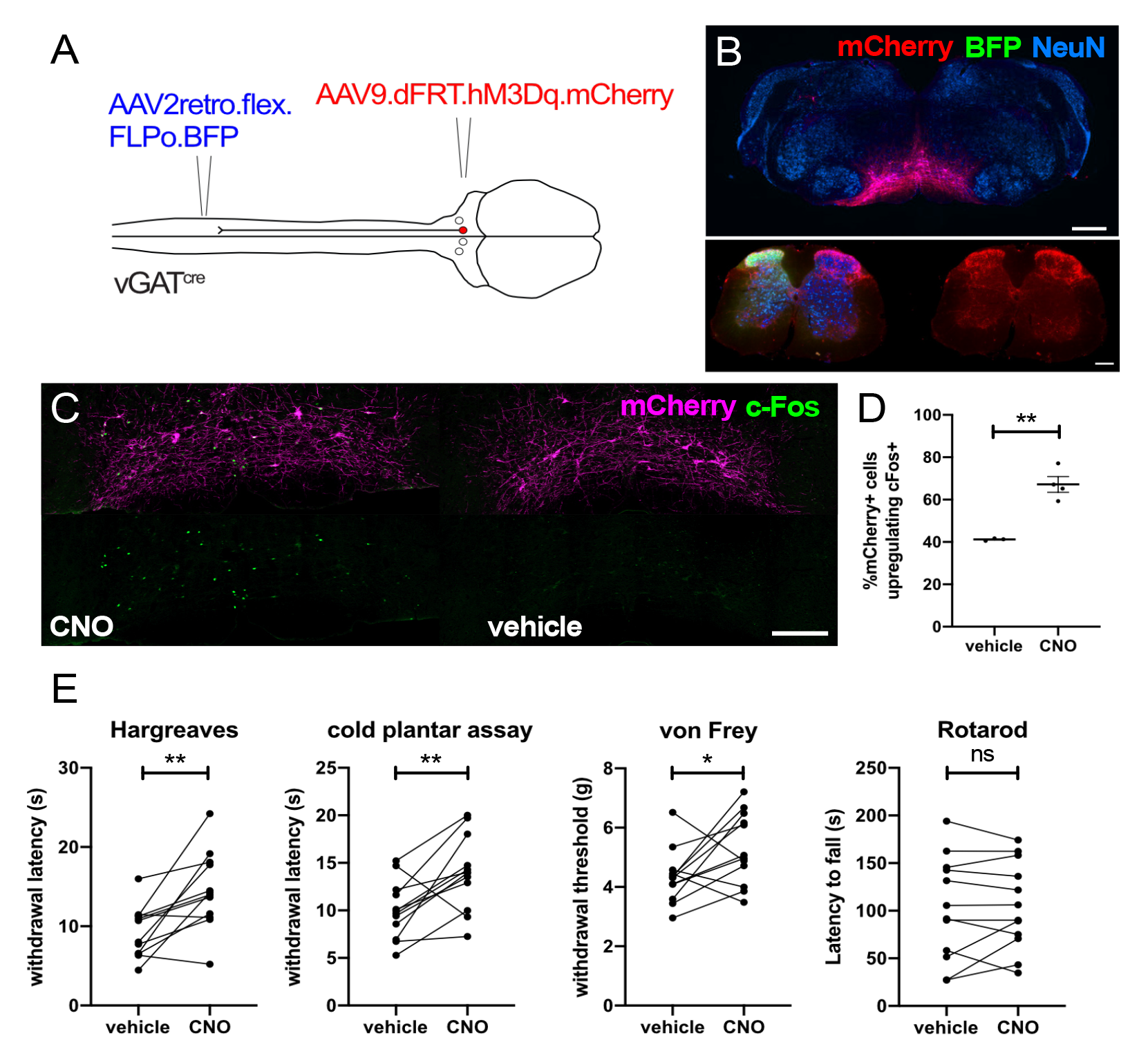
Chemogenetic activation of descending vGAT^cre^ RVM neurons inhibits cutaneous sensitivity: A. Injection scheme for introducing hM3Dq to vGAT^cre^ descending RVM projections to the left lumbar spinal cord. B. Injection sites from the hindbrain and lumbar spinal cord of a representative animal, note the presence of axons on both injected and non-injected sides of the spinal cord (scale bar = 200 μm for hindbrain and 50 μm for the spinal cord image). C. images of c-Fos in hM3Dq-mCherry expressing RVM neurons following either vehicle or CNO injection (scale bar = 100 μm). D. Quantification of mCherry+ cells that upregulate c-Fos after CNO injection (vehicle (n = 3) vs CNO (n = 4) p = 0.002 unpaired t-test). E. Activation of vGAT^cre^ descending RVM neurons with CNO increased withdrawal latencies to thermal stimuli and von Frey thresholds when compared to vehicle injected controls. (Hargreaves, vehicle vs CNO p = 0.0011; cold plantar assay, vehicle vs CNO p = 0.0089; von Frey vehicle vs CNO p = 0.0045, paired t-tests). Sensorimotor coordination was unaffected by activation of vGAT^cre^ RVM PNs (Rotorod, Vehicle vs CNO p = 0.699). N = 12 animals for all tests. Significance levels; * p < 0.05, ** p < 0.01.

### Inhibitory RVM projections form functional synapses throughout the spinal cord and can mediate widespread thermal antinociception

The presence of wide-ranging axon collaterals in multiple spinal segments raises the possibility that these neurons control nociception widely, not only on both side of the body but also extending significantly along the rostrocaudal axis. To address this question, we tested whether RVM neurons labelled from a single unilateral lumbar spinal cord injection site would form functional synapses along the spinal cord.

To test for the presence of functional synapses in multiple spinal segments, we expressed ChR2 in vGAT^cre^ neurons that projected to the lumbar spinal cord using the intersectional strategy (Figure 4A). We then prepared transverse spinal cord slices from both the lumbar and cervical segments and performed the optogenetic stimulation experiment. This strategy produced the expected labelling of hindbrain neurons, with ChR2-YFP being restricted to the projection neurons retrogradely labelled with mCherry (Figure 4A). leIPSCs were recorded from neurons in both lumbar and cervical spinal slices, which were blocked by bicuculline and strychnine (average leIPSC amplitude in cervical neurons 611.9 ± 312 pA, and in lumbar neurons 191.8 ± 63.4 pA) (Figure 4B and C). This demonstrates that the axon collaterals of descending RVM neurons form functional synapses within anatomically distinct sites, potentially capable of producing wide-ranging inhibition throughout the spinal cord.

**Figure 4:**
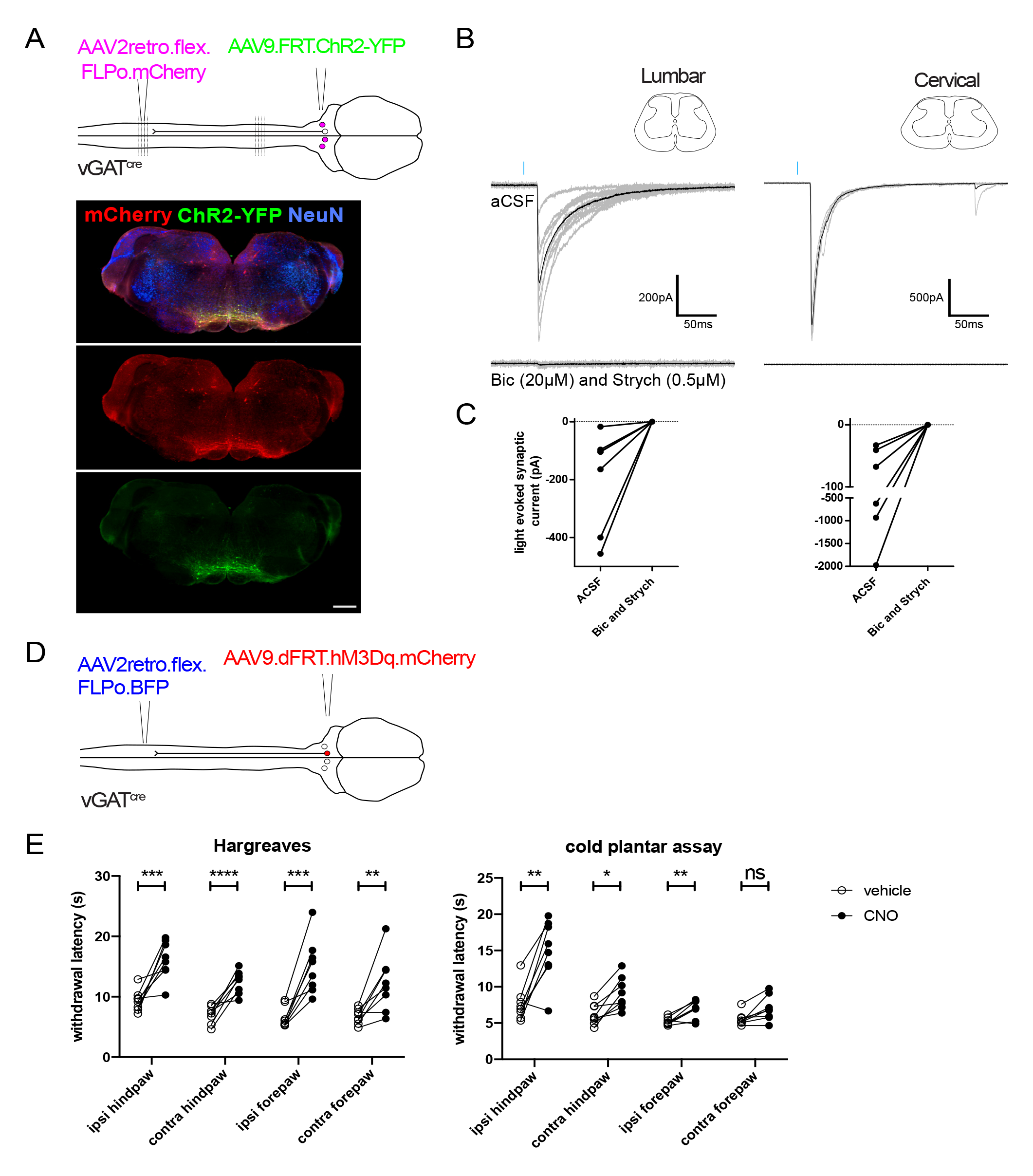
vGAT^cre^ RVM neurons form functional synapses throughout the spinal cord and can globally reduce thermal sensitivity. A. Intersectional strategy for expressing ChR2 in vGAT^cre^ neurons that project to the lumbar spinal cord. Injection site in the brainstem showing projection neurons retrogradely labelled with mCherry that were intersectionally labelled with ChR2-YFP, scale bar = 200 μm. B. Example light-evoked synaptic currents taken from neurons recorded in lumbar and cervical spinal cord slices from a vGAT^cre^ animal that received the injections depicted in A. Averaged traces from 10 stimuli are shown in black, and the individual traces are shown in grey. Both light-evoked currents are completely blocked by bicuculline and strychnine application. C. Group data from neurons recorded in slices of cervical or lumbar spinal cord that received leIPSCs following blue light stimulation (cervical n = 6, lumbar n = 7). D. Intersectional strategy for expressing hM3Dq in descending inhibitory neurons that project to the lumbar spinal cord. E. Withdrawal latencies from all four paws to heat and cold stimuli following CNO or vehicle injection. Response latencies were significantly increased in all paws for the Hargreaves assay (ipsi hindpaw p = 0.000328, contra hindpaw p = 0.000072, ipsi forepaw p = 0.000492, contra forepaw p = 0.006408, adjusted p values), and in 3 out of 4 paws for the Cold plantar assay (ipsi hindpaw p = 0.003188, contra hindpaw p = 0.016075, ipsi forepaw p = 0.009658, contra forepaw p = 0.051301, adjusted p values) (repeated t-tests with adjusted p values, Bonferroni-Dunn post hoc correction). N = 8 animals, *p < 0.05, **p < 0.01, ***p < 0.001, ****p < 0.0001.

The results of the optogenetic experiments together with the anatomical tracing suggests that inhibitory RVM projection neurons could include a population that globally inhibits nociception. We therefore repeated our chemogenetic experiments with unilateral lumbar AAV injections and tested whether CNO injection would prolong thermal withdrawal latencies not only in the hindpaws but also in the forepaws (Figure 4E). We found that the response latencies for the forepaws were also increased to thermal stimuli, and as expected from the bilateral anatomic tracing experiments, we also observed increases in the contralateral paws, although those were generally smaller than those on the ipsilateral side (average difference in latency between vehicle and CNO for Hargreaves 6.79, 5.39, 8.43, and 5.49 s for the ipsilateral hindpaw, contralateral hindpaw, ipsilateral forepaw, and contralateral forepaw respectively). Similar but smaller effects were seen in the cold plantar assay, where the magnitude of the effect was smallest for the forepaw contralateral to the spinal injection site (average difference in latency between vehicle and CNO = 7.33, 2.88, 1.71, and 1.46 s for the ipsilateral hindpaw, contralateral hindpaw, ipsilateral forepaw, and contralateral forepaw respectively). Therefore, descending inhibitory projections to the lumbar spinal cord terminate in multiple spinal segments and can reduce thermal sensitivity in multiple areas.

### Tonic activity of descending vGAT^cre^ projection neurons limits mechanical hypersensitivity, tactile allodynia, and spontaneous pain

Chemogenetic experiments demonstrated that activation of descending RVM inhibitory neurons reduces sensitivity to cutaneous stimuli, but whether these neurons also regulate baseline sensitivity remains to be explored. To test whether these neurons determine baseline nociception, we complemented our gain-of-function with loss-of-function experiments. For this, we injected the lumbar spinal cords of vGAT^cre^ mice or cre negative littermates unilaterally with AAV2retro containing a Cre-dependent Flpo. One week later, the hindbrains of these same animals were injected with AAV containing a Flpo-dependent tetanus toxin light chain (TetxLC) coding sequence to block neurotransmitter release from the descending inhibitory neurons (Figure 5A and B)^25^. Within seven days of hindbrain injection of AAV9.dFRT.TetLC.eGFP, Cre-expressing animals developed strong hypersensitivity to both punctate and dynamic mechanical stimuli, hallmarks of mechanical allodynia, (von Frey thresholds: pre-injection: 3.48 ± 0.2, post-injection: = 1.92 ± 0.3 g; tactile allodynia scores: pre-injection: 0.7 ± 0.1, post-injection: 1.84 ± 0.2) (Figure 5C). Strikingly, these animals also exhibited spontaneous aversive behaviors, such as flinching of the ipsilateral hindpaw, and a reluctance to use this paw for locomotion (video 2). Gross sensorimotor coordination assessed in the rotarod test remained unaltered (Figure 5C). In contrast to the chemogenetic activation experiments, silencing the vGAT^cre^ descending neurons did not significantly alter thermal sensitivity, suggesting that these neurons are not involved in the control of heat and cold sensitivity under resting conditions (Figure 5C). They might however be recruited in certain conditions such as stress or chronic pain. Although the observed changes largely affected the injected side of the spinal cord, we also observed an increase in sensitivity to dynamic mechanical stimuli on the contralateral paw (tactile allodynia score; pre injection: 0.69, post injection: 1.26) (Figure 5C).

**Figure 5:**
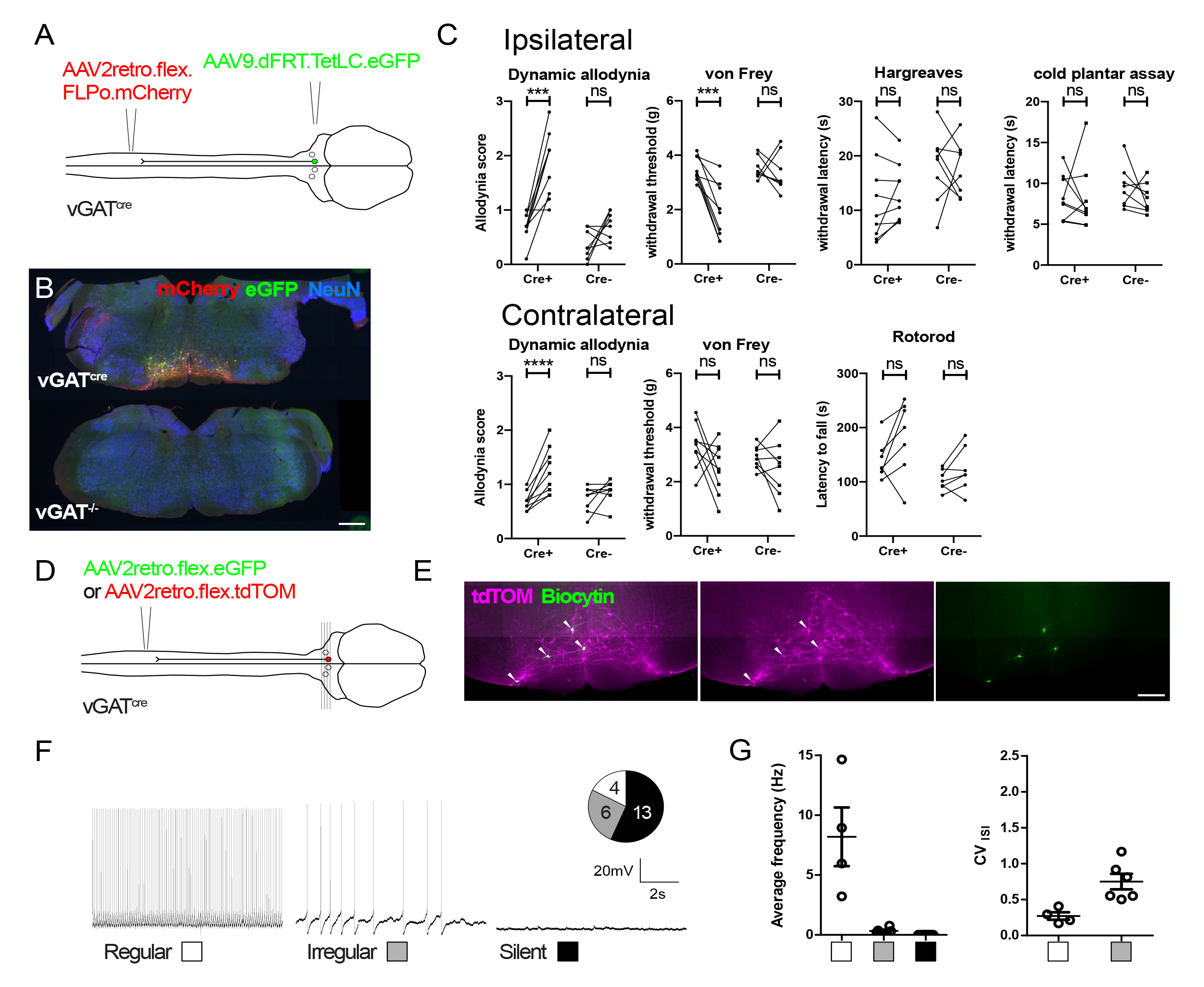
Tetanus toxin-mediated silencing of descending vGAT^cre^ RVM neurons produces mechanical allodynia. A. Injection scheme for silencing vGAT^cre^ PNs in the RVM with TetxLC. B. Example injection sites in the hindbrain from vGAT^cre^ mice and Cre-littermates, illustrating the expression of mCherry and eGFP in the same neurons, and both proteins only being present in the hindbrains of vGAT^cre^ animals (scale bar = 100 μm). C. Following the hindbrain injection, Cre-expressing animals were more sensitive to von Frey punctate stimulation and dynamic brush stimulation (von Frey F(1,15) = 9.6, p = 0.0071, for Cre+ p = 0.0002, for Cre-p > 0.99, dynamic allodynia test F(1,15) = 8.9, p = 0.0091, Cre+ p < 0.0001, Cre-p = 0.212,); whereas responses to thermal stimuli were unaltered. (Hargreaves F(1,15) = 0.3, p = 0.59 for Cre+ p > 0.99, for Cre-p > 0.99; cold plantar assay F(1,15) = 0.3 p = 0.54, Cre+ p > 0.99, Cre-p = 0.687,). An increase in dynamic allodynia is also seen in the contralateral paw (F(1,15) = 8.6, p = 0.010, Cre+ p < 0.0001, Cre-p = 0.334), whereas punctate von Frey thresholds were not significantly reduced (von Frey F(1,15) = 0.54 p = 0.47, Cre+ p < 0.0001, Cre-p = 0.334). No alterations in gross sensorimotor coordination were detected in the Rotarod assay (F(1,12) = 0.8, p = 0.37, Cre+ p = 0.066, Cre-p = 0.60). All comparisons in behavioral assays are pre vs post RVM injection using a 2-Way ANOVA with Bonferonni post hoc tests. D. Injection scheme for labelling descending vGAT^cre^ projection neurons for targeted whole-cell recordings. E. Hindbrain slice containing tdTomato+ cells that were labelled with biocytin during whole-cell recordings (scale bar = 100 μm). F. Representative traces from vGAT^cre^ RVM projection neurons recorded in current clamp mode at 0 pA that could be assigned to different groups based on their spontaneous activity. Inset illustrates the frequency at which these different cell types were observed. G. Average frequency of spontaneous action potential firing for each cell type, and the coefficient of variance for the interspike intervals (CV ISI) of irregular and regular firing vGAT^cre^ projection neurons. Regular neurons have a consistently lower CV ISI than irregular neurons indicating less variability in the time between action potentials.

These data could suggest that descending vGAT^cre^ neurons include populations whose activation can inhibit sensitivity to varied stimuli, and other populations that are continuously active under normal circumstances and limit mechanical sensitivity. Accordingly, there must be neurons within the vGAT^cre^ descending population with lower basal activity that can be recruited to reduce cutaneous, including thermal, sensitivity, and others that have ongoing activity to prevent mechanical allodynia. We performed targeted whole-cell recordings from labelled RVM vGAT^cre^ descending neurons to test whether these activity patterns existed (Figure 5 D-G). In agreement with this prediction, recorded neurons could be broadly divided into three categories based on their spontaneous action potential firing patterns ^26^. These included spontaneously active neurons with regular and irregular firing as well as inactive cells (referred to as regular, irregular and silent respectively) (Figure 5F), consistent with the findings of others^26^). Regular and irregular firing neurons could be distinguished from each other based on the coefficient of variance of their interspike intervals (Figure 5G)^27^. Regular firing neurons exhibited a wide range of firing frequencies (3.2 – 14.7 Hz), which were consistently higher than the irregular firing neurons (0.13 – 0.77 Hz) (Figure 5G). Collectively this indicates that descending inhibitory RVM neurons are a functionally heterogeneous population, including cells capable of reducing cutaneous sensitivity to thermal and mechanical stimuli and others required for limiting sensitivity to mechanical stimuli and preventing allodynia.

## Discussion

Descending pain modulation is an adaptive system for controlling pain perception in a context-dependent manner. This is important for managing pain in a variety of situations and emotional states, in order to respond appropriately to threats, environmental changes or damaging stimuli ^28^. In addition, components of these descending systems are found to be altered in many forms of chronic pain ^29, 30^. Identifying the components of this system is therefore essential to understand the processes that regulate pain sensitivity and to this end, we have characterized a broad population of inhibitory neurons in the RVM that are capable of suppressing sensitivity to external stimuli at the level of the spinal cord. Subsets of the RVM inhibitory projection neurons have been identified and functionally interrogated in previous work, but the detailed anatomy of the projections is less well characterised^11, 23, 24^. In general, most of the neurons in the RVM that we could retrogradely label from the spinal cord with AAV2retro contained either vGAT or vGluT2 mRNA. However, we found that inhibitory neurons were a small proportion of the total population of descending projection neurons in the ventral hindbrain, in agreement with recent transcriptional profiling studies^31^. However, these studies may not include all projections due to the known tropism of AAV2retro vectors^12, 32^. We decided to study the inhibitory population, although there are almost certainly other functionally important groups of projection neurons in this area, many of which were not captured in our sample, such as the serotonergic neurons of the raphe magnus ^13, 33, 34^.

### Anatomical organization of descending inhibitory RVM projections

Using an intersectional approach, we were able to selectively label and manipulate inhibitory projection neurons in the RVM that innervate the lumbar spinal cord. Strikingly, this population had wide-ranging axonal projections that terminated within multiple somatotopically distinct spinal segments. These anatomical features would make them ideally placed for global pain modulation, for example during stress-induced analgesia or other forms of diffuse noxious inhibitory control ^35, 36^. Indeed, when activated, the inhibitory population was capable of rather global reductions in diverse modalities of cutaneous sensitivity, such as to cold and heat stimuli. However, the extent of this reduction between somatotopic regions differed, with the more separated somatotopic areas having smaller decreases in sensitivity to heat and cold compared to those innervating the spinal segment from which the projections were retrolabelled. Although there are clearly RVM neurons with wide-ranging axons throughout the length of the spinal cord, it is uncertain that all descending neurons contribute to this, and there is potentially some regional specificity of these projections. This agrees with the retrograde labelling from the spinal cord preferentially labelling RVM neurons on the ipsilateral side ^37^. This labelling in the hindbrain could either be due to some RVM neurons innervating specific spinal segments or sides, or that the axonal innervation from many RVM neurons is global, but with certain regions being preferentially innervated relative to others. Labelling of individual RVM projection neurons is required to address these questions.

Regional specificity of altered sensitivity was also seen in the loss-of-function experiments. When vGAT^cre^ projection neurons were silenced, there was a smaller hypersensitivity to dynamic tactile stimuli measured on the contralateral paw, suggesting that these neurons may also be required for widespread tuning of normal sensitivity. However, this was not detected in tests of punctate mechanical sensitivity. Further, in some central sensitization syndromes, such as fibromyalgia and chronic regional pain syndrome, widespread pain hypersensitivity is possibly the consequence of pathological changes within the pain processing centers of the CNS ^38–40^. With widespread projections throughout the rostrocaudal extension of the spinal cord, together with their strong tonic influence on normal tactile sensitivity, it is tempting to speculate that these inhibitory RVM projection neurons or subsets of them could be involved in such conditions.

Intriguingly, these spinally projecting neurons also contained branches that innervated other CNS regions, including the spinal trigeminal nucleus, LPb, and PAG. All these areas are sensory structures, either receiving nociceptive information from peripheral tissues, or areas where ascending nociceptive information is relayed from the spinal cord ^19–21, 41, 42^. It is therefore possible that these neurons can influence the processing of sensory information at supraspinal sites, potentially regulating the aversive and affective dimensions of sensory information ^43–45^.

### Functional roles of descending vGAT^cre^ RVM neurons

Subsets of inhibitory RVM projection neurons have been characterized in previous studies, such as those expressing Kappa opioid receptors (KOR) and pro-enkephalin (pENK) ^23, 24^. In agreement with our findings, activation of these groups generally has an inhibitory effect on most nociceptive modalities^24^. Similarly, studies have shown that inhibiting the vGAT^cre^ descending projections has little influence over thermal sensitivity, both with chemogenetic and optogenetic approaches ^11^. In contrast to our findings, tetanus-mediated silencing of pENK-or GABA-containing hindbrain neurons increased sensitivity to thermal stimuli without affecting mechanical sensitivity ^23^. Additionally, other studies demonstrate that chemogenetic inhibition of the RVM inhibitory neurons reduces sensitivity to mechanical stimuli through disinhibition of spinal inhibitory neurons, whereas in the present study, strong mechanical hypersensitivity and spontaneous pain-like responses were observed^11^. These differences between studies are likely explained by differences in the neuronal populations that were targeted in the hindbrain and the efficacy of the various strategies for inhibiting, silencing, and activating these neurons (chemogenetics and optogenetics vs tetanus toxin-mediated silencing). In our approach, we limited the expression of viral transgenes to the spinally projecting inhibitory neurons, established a method of targeting as many retrogradely traced neurons as possible, and used a method to permanently silence the neurons. This approach resulted in strong inhibition of most inhibitory neurons that can be retrogradely traced from the spinal cord with AAV vectors, and therefore may produce phenotypes that differ from those seen by others. Additionally, we found that the medullary projection neurons are heterogeneous in terms of their projection targets, electrophysiological properties and anatomical location. Therefore, it is highly likely that targeting different neuron groups within this larger population will have diverse effects on the sensory system, which are not revealed by silencing or activating the broader population. Further studies are required to confirm these suppositions.

The data here provide evidence that inhibitory projections neurons of the hindbrain that project their axons to the spinal dorsal horn are critical components controlling physiological pain sensitivity and are capable of powerful suppression of nociception when activated. This potentially reflects functional heterogeneity within this population, and could be an important dysfunctional element in widespread pain conditions. These results provide a starting point for further dissection of these descending pathways and highlights the importance of descending inhibitory projections of the RVM for regulating basal sensitivity to external stimuli.

## Acknowledgements

We would like to thank the Center for Microscopy and Image Analysis at the University of Zurich for assistance with the tissue clearing and light-sheet microscopy experiments. We would also like to thank Louis Scheurer, Isabelle Kellenberger and Katharina Struckmeyer-Fichtel for technical assistance.

## Methods

### Animals

Mice aged between 6-12 weeks were used for behavioral and anatomical labelling experiments. For electrophysiology experiments, animals aged between 3-4 weeks were injected and were prepared for slice recordings >10 days later. Permission to perform these experiments was obtained from the Veterinäramt des Kantons Zürich (154/2018 and 097/2021). All transgenic mouse lines used in this study are listed in the key resources table.

### Surgeries

The dorsal horn of the lumbar spinal cord was injected as previously reported elsewhere ^46, 47^. Mice were anesthetized with 5% Isoflurane which was maintained at 1-3% through a face mask during all surgeries, analgesia (buprenorphine subcutaneously injected at 0.1-0.2 mg/kg) was provided before and after each surgery. During surgery the body temperature was maintained using a heated mat and vitamin A cream was applied to protect the eyes.

The skin above the injection site was shaved and disinfected with Betadine, and an incision was made to expose the T13 vertebra. The surrounding tissues were separated from the vertebral column and the T13 vertebra was clamped in position using spinal adaptors. A borehole was made in the middle of the T13 vertebra on one side and AAVs were injected into the spinal dorsal horn at a depth of 200-300 μm below the spinal surface approximately 500 μm lateral to the central artery. In some anatomical tracing experiments both sides of the spinal cord were injected in a similar manner. For virus injections, 3 x 300 nl virus solution was injected along the rostrocaudal extent of the spinal cord at an infusion rate of 50 nl/min. The viral vectors used in this study can be found in the key resources table.

For injections into the hindbrain the head was fixed in position using adjustable cheek bars. Injection coordinates were determined by the location of the retrogradely labelled hindbrain neurons that were identified in initial experiments, which relative to bregma were, and −5.8, ±0.5, 5.9 (rostrocaudal, mediolateral, dorsoventral respectively). Viruses were infused at 50- 100 nl/min to a total volume of 1 μl for each injection. Injections were made using a motorized frame controlled by Neurostar Stereodrive software. The skull was installed in the frame to be as straight and as flat as possible, (difference in z position between lambda and bregma < 100 μm). To compensate for variations in tilt and scaling, adjustments to the injection target were made in the software relative to four positions on the surface of the skull (Bregma, Lambda, 2 mm to the right of the midline, and 2 mm to the left of the midline).

### General features of tissue preparation and immunohistochemistry

Animals were perfusion fixed with freshly prepared 4% paraformaldehyde (PFA) (room temperature, dissolved in 0.1 M PB, adjusted to pH 7.4). Transcardial perfusion with fixative followed the removal of blood the mouse circulatory system by perfusing with 0.1 M PB. Nervous tissues were quickly dissected and post-fixed in 4% PFA at 4°C for two hours. For CLARITY tissue clearing experiments, tissues were post-fixed for 24 hours after dissection (see below for more details). After post-fixation, brains or spinal cords were placed in 30% sucrose solution (dissolved in 0.1 M PB) for 24-72 hours for cryoprotection of the tissue.

These were then rinsed with 0.1 M PB before being embedded in NEG-50 mounting medium. Tissues were sliced at 60 μm on a sliding blade microtome (Hyrax KS 34, histocam AG) and stored as free-floating sections. For long-term storage, free-floating sections were stored in antifreeze medium (50 mM sodium phosphate buffer, 30% ethylene glycol, 15% glucose, and sodium azide (200 mg/L) at −20°C until required.

For immunostaining, sections were first washed three times in 0.1M PB and incubated in 50% EtOH for 30 minutes at room temperature, followed by three rinses in PBS with added salt (8 g NaCl per liter of PBS). Sections were incubated in a mixture of primary antibodies for two nights at 4°C. The antibodies were reconstituted in a solution containing PBS with added salt, 0.03% Triton-X, and 10% normal donkey serum. All primary antibodies and dilutions used in this study are listed in the key resources table. Tissue sections were rinsed 3 times in PBS with added salt before being incubated in species specific secondary antibodies overnight at 4°C (see key resources table). Finally, tissue sections were rinsed three times in PBS with added salt and mounted on microscope slides with Dako anti-fade medium.

### Tissue clearing and light sheet microscopy

Following 24 hour post-fixation in 4% PFA at 4°C, tissues were placed in a pre-chilled hydrogel solution (containing 45 ml PBS, 5 ml 40% polyacrylamide, and 125mg VA-044) sealed in an air tight container for >2 days at 4°C. Samples were baked at 37°C for 3 hours to allow polymerization of the hydrogel. Tissues were then transferred to an SDS-based clearing solution (containing 24.7g boric acid, 80g Sodium dodecyl sulfate in 2L ddH_2_O, adjusted to pH 8.5), and lipids were removed by active electrophoretic clearing for 3-5 hours^48^. The clearing solution was circulated and chilled throughout the tissue clearing to avoid SDS warming and precipitation. Following active clearing, tissues were rinsed twice in PBS-T (containing 0.1% Triton-X) and then placed in histodenz solution overnight to allow for refractive index matching (refractive index adjusted to 1.4655 with PBS). Samples were glued to a platform and placed in an imaging cuvette filled with index-matched histodenz before being installed in a MesoSPIM light sheet microscope^22^. Images were acquired using MesoSPIM control and processed with ImageJ/FIJI.

### RNA scope fluorescent in situ hybridization (fISH)

For fluorescent multiplex *in situ* hybridization experiments, brains were quickly dissected following euthanasia (< 5 minutes). The hindbrains were isolated from the forebrain and cerebellum before being rapidly frozen in liquid nitrogen. Hindbrain tissues were embedded in Neg50 mounting media which was frozen for cryostat sectioning.

Tissue blocks were installed in a cryostat (Hyrax 60, histocam AG), and sections were taken at 20 μm thickness which were mounted directly onto superfrost microscope slices. Slides were stored at −80°C before fISH experiments. All *in situ* hybridization experiments followed the guidelines according to the RNAscope® Fluorescent Multiplex *in situ* hybridization v1 kit. Tissues were fixed for 15 min at 4°C in freshly prepared 4% PFA dissolved in PBS. Following fixation, the slides were dehydrated through an alcohol series (50%, 70%, 100% and 100% again), for 5 minutes at each EtOH concentration. The tissue containing region of the slide was delineated with a heat resistant fat pen and allowed to dry. Tissues were treated with a Protease solution (Protease IV) for 30 min at room temperature before hybridization. Protease was removed and slides were briefly rinsed twice in PBS, Z probes (ACD bio) were hybridized to the tissues for 2 hours in a 40 °C purpose build oven. Probes were removed and slides were rinsed twice in wash buffer for 2 minutes each before and between all amplification stages. Amplification of the hybridized probes were achieved using the amplification reagents 1, 2, 3, and finally were either labelled with 4A, 4B, or 4C (for 30, 15, 30, and 15 minutes respectively) in a 40°C oven. Slides were counterstained with DAPI for 10 minutes and mounted in DAKO antifade mounting medium. The probes used in this study are listed in the key resources table.

Cells were counted using a CellProfiler pipeline to identify nuclei with DAPI staining, and to assign fluorescent puncta from each detected mRNA species within a small area surrounding the nuclei. Cells were counted as positive if they contained above the 90^th^ percentile of puncta/nuclei for each fluorophore, determined by running a 3-plex negative control on an adjacent slide in parallel with the same amplification reagents ^49^.

### Image acquisition and analysis

For an overview of injection sites and imaging entire brain sections, epifluorescence images were acquired using a Zeiss Axio Scan.Z1 slidescanner. For quantification of retrogradely labelled cells in the hindbrain, image stacks were acquired at 5 μm z-spacing using a Zeiss LSM 800 confocal microscope. Confocal scans were made using 488, 561 and 640 nm lasers and the pinhole was set to 1 Airy Unit for reliable optical sectioning. Image stacks were acquired where immunoreactivity of all antigens was clearly visible. Acquisition settings were kept the same within each quantification experiment and acquired images were analyzed offline with FIJI using the cell counter plugin. Data were processed in Microsoft Excel and were presented and analyzed with GraphPad Prism 8.

### Behavioral assays

For activation or silencing of descending RVM projections, we used an intersectional strategy in which an AAV2retro.flex.FLPo virus (also containing either a BFP or mCherry coding sequence) was used to transduce descending Cre-expressing neurons. One-week later mice received hindbrain injections of AAVs containing a FLPo-dependent effector (either hM3Dq or TetLC). For DREADD experiments, 10 days incubation time was given to allow the expression of the receptor, whereas 5-7 days were given for TetxLC expression before sensory and motor testing. Before experiments mice were acclimatized to the behavioral setup for at least one hour. For the Hargreaves plantar, cold plantar, electronic von Frey, and Rotarod assays, an average from six measurements was recorded. For tactile allodynia assays, an average from 10 response scores was recorded. All measurements were taken from both hindlimbs of all animals, and in some experiments, the forepaws were also tested, using the same scoring criteria for testing hindpaws.

### Hargreaves

To measure heat sensitivity, paws were stimulated with an infrared heat source (Hargreaves plantar assay IITC). Mice were placed on a transparent platform preheated to 30°C, and withdrawal latencies were recorded in response to heating from an infrared heat source using an inbuilt timer. The stimulation intensity was set to 20% heater power for each test, and a cutoff time of 32 s was used to prevent potential tissue damage, with 3-5 min between stimulating the same paw to avoid tissue sensitization.

### Cold plantar assay

Mice were placed on a 5 mm borosilicate glass platform and were stimulated from beneath with dry ice pellets. The time taken between touching the glass from below to the withdrawal of the paw was measured and 20 s was used as the cutoff time to avoid any tissue damage. The same paw was only tested again after a 3-5 min recovery time.

### Electronic von Frey

Mechanical thresholds were determined using an electronic von Frey algesiometer (IITC). Animals were adapted on a mesh surface, and the plantar surface of each paw was stimulated with a bendable plastic filament attached to a pressure sensitive probe. Pressure was applied to the plantar surface gradually at a constant speed until the animal withdrew its paw. The maximum pressure that was present when the animal withdrew its paw was recorded.

### Dynamic allodynia test

Sensitivity to dynamic mechanical stimuli was a measured by gently brushing the hindpaw plantar surface with a paintbrush at a constant speed from heel to toe. Allodynia/Brush response scores were given for each stimulation were based on methods previously described, with some adaptations to the scoring ^50, 51^. In brief, animals that did not withdraw their paw were given a score or 0, and animals that quickly withdrew were given a score of 1. Animals that withdrew and kept the paw raised (extended lifting) for > 0.5 s were given a score of 2, and animals that flinched or licked the paw post-stimulation were scored as 3. Repeated flinching and licking of the stimulated paw was given a score of 4.

### Rotarod

Sensorimotor coordination was evaluated using an accelerating rotarod, and the time taken for animals to fall from the rotating barrel was recorded. The barrel rotated from 4 - 40rpm accelerating at a constant rate over a period of 300 st. Values were not included if the animal jumped from the barrel in the direction of rotation, and if the animal jumped in >50% of trials for a given time point these data were discarded.

### Drug application

Clozapine-N-oxide (CNO) was injected at a concentration of 2 mg/kg intraperitoneally for DREADD-mediated neuronal activation experiments. Animals were tested directly before and 1 – 3 hours after i.p. injections. Stock CNO was stored at a 500x concentration in DMSO at 100 mg/ml, which was diluted in sterile filtered saline immediately prior to injection.

### Slice preparation, optogenetics and whole cell recording

Spinal cord slices were prepared for electrophysiology experiments in a similar manner to previous studies ^52^. Mice were decapitated while under isoflurane anesthesia, and the vertebral column was rapidly dissected and placed in oxygenated ice-cold dissection solution (containing in mM: 65 NaCl, 105 sucrose, 2.5 KCl, 1.25 NaH_2_PO_4_, 25 NaHCO_3_, 25 glucose, 0.5 CaCl_2_, 7 MgCl_2_). The back of the animal was pinned to the bottom of a sylgard-coated plate with the ventral surface facing up, and a laminectomy was performed to expose the spinal cord. The spinal cord was then carefully removed from the vertebral column and the lumbar and cervical enlargements were isolated and sliced in the transverse plane. The ventral side of each spinal cord enlargement was then glued to an agar block, mounted in a slicing chamber and installed in a vibrating blade microtome (D.S.K microsclicer DTK1000). Transverse slices of 300 – 350 µm thickness were taken and transferred to a holding chamber filled with oxygenated aCSF (containing in mM: 120 NaCl, 26 NaHCO_3_, 1.25 NaH_2_PO_4_, 2.5 KCl, 5 HEPES, 14.6 glucose, 2 CaCl_2_, 1 MgCl_2_; pH 7.35 - 7.40, osmolarity 305 - 315 mOsm) warmed to 34°C. Slices were left to recover for at least 30 min before recordings were taken.

For the preparation of hindbrain slices, brains were quickly removed following euthanasia of the animal. The brain was placed in ice cold dissection solution, and the hindbrain was isolated by removing the forebrain and cerebellum. The hindbrain was glued to an agar block and transverse sections were cut at 250 µm, which were held in an incubation chamber filled with warm aCSF (34°C) until recordings were taken.

Targeted whole-cell recordings were taken at room temperature using a HEKA EPC10 amplifier with Patchmaster software (HEKA Elektronik) at a sampling frequency of 20kHz. For optogenetic experiments, cells in the spinal dorsal horn (laminae I-III) were randomly targeted using DIC optics. A cesium-based internal solution was used (containing in mM: 120 CsCl, 10 HEPES, 0.05 EGTA, 2 MgCl_2_, 2 Mg-ATP, 0.1 Na-GTP, 5 QX-314) and recordings were taken in voltage clamp mode, with blue light stimuli (4ms, 473nm) delivered through the objective lens at a frequency of 0.1 Hz from a monochromator (TILL photonics). To confirm the inhibitory nature of light evoked synaptic events, bicuculline (20 µM) and strychnine (0.5 µM) were bath applied to the slice during recording.

For targeted recordings of RVM projection neurons, the spinal cords of vGAT^cre^ animals were injected with AAV2retro.flex.tdTOM or AAV2retro.flex.eGFP one week before the preparation of hindbrain slices. Labelled cells were visualised by epifluorescent illumination using a monochromator and were targeted for whole-cell recording. Recordings were taken in current clamp mode using a potassium gluconate internal solution (containing in mM, 130 K-Gluconate, 5 NaCl, 1 EGTA, 10 HEPES, 5 Mg-ATP, 0.5 Na-GTP, 2 biocytin) to determine spontaneous activity. Following recording, the recording electrode was slowly retracted, and the slice was fixed overnight at 4° in 4% PFA. Fixed hindbrain slices were processed to reveal biocytin-labelled cells and immunoreacted to confirm their tdTOM or eGFP content.

Access resistance was monitored throughout each experiment, and traces from each experiment were analyzed further if the access resistance changed <30% during the recording.

### Experimental design and statistical tests

All chemogenic experiments were performed twice on all animals so that each received both CNO and vehicle on different experimental days, with the experimenter being blinded to the substance injected. Comparisons were made between CNO or vehicle treated animals. Statistical significance was taken as *p* < 0.05. For behavioral experiments involving tetanus toxin-mediated silencing of neurons, littermates that did not express Cre were used as controls and the experimenter being blinded to the genotype. Comparisons were made before and 5-7 days after the hindbrain injection of AAVs containing the coding sequence for tetanus toxin light chain using a 2-Way ANOVA to determine the interaction of genotype and time.

### Data collection, storage, and presentation

Data were analyzed and presented using GraphPad Prism 8. Figures were arranged in Affinity Designer 2. Data are available on reasonable request and are kept on the internal servers of the Institute for Pharmacology and Toxicology, University of Zürich.

| **Antibody** | **Supplier/Source** | **Cat#/RRID** |
| --- | --- | --- |
| Chicken anti-GFP (1:1000) | LifeTech | A10262/<br>AB_2619988 |
| Goat anti-mCherry (1:500) | Sicgen | AB0081-200/AB_2333094 |
| Guinea pig-NeuN (1:1000) | Synaptic systems | 266004/AB_2619988 |
| Rabbit anti-c-Fos (1:500) | Calbiochem | PC38/AB_2106755 |
| Alexa 488-Donkey anti-Chicken (1:500) | Jackson ImmunoResearch | 703-546-155/AB_2340376 |
| Cy3-Donkey anti-goat (1:500) | Jackson ImmunoResearch | 705-166-147/AB_2340413 |
| Alexa 488-Donkey anti-goat (1:500) | Jackson ImmunoResearch | 795-546-147/ - |
| Alexa 647-Donkey anti-guinea pig (1:500) | Jackson ImmunoResearch | 706-496-148/ - |
| Alexa 647-Donkey anti-rabbit (1:500) | Jackson ImmunoResearch | 711-607-003/AB_2340626 |
| Cy3-Donkey anti-rabbit (1:500) | Jackson ImmunoResearch | 711-165-152/AB_2307443 |

| **Target** | **Supplier/Source** | **Cat#** |
| --- | --- | --- |
| EGFP | Advanced Cell Diagnostics | 400281 |
| EGFP-C3 | Advanced Cell Diagnostics | 400281-C3 |
| Mm-Slc17a6 | Advanced Cell Diagnostics | 319171 |
| Mm-Slc17a6-C2 | Advanced Cell Diagnostics | 319171-C2 |
| Mm-Slc32a1 | Advanced Cell Diagnostics | 319191 |
| Mm-Slc32a1-C2 | Advanced Cell Diagnostics | 319191-C2 |
| 3-plex negative control probe | Advanced Cell Diagnostics | 320871 |
| RNAscope Fluorescent Multiplex Reagent Kit | Advanced Cell Diagnostics | 320850 |

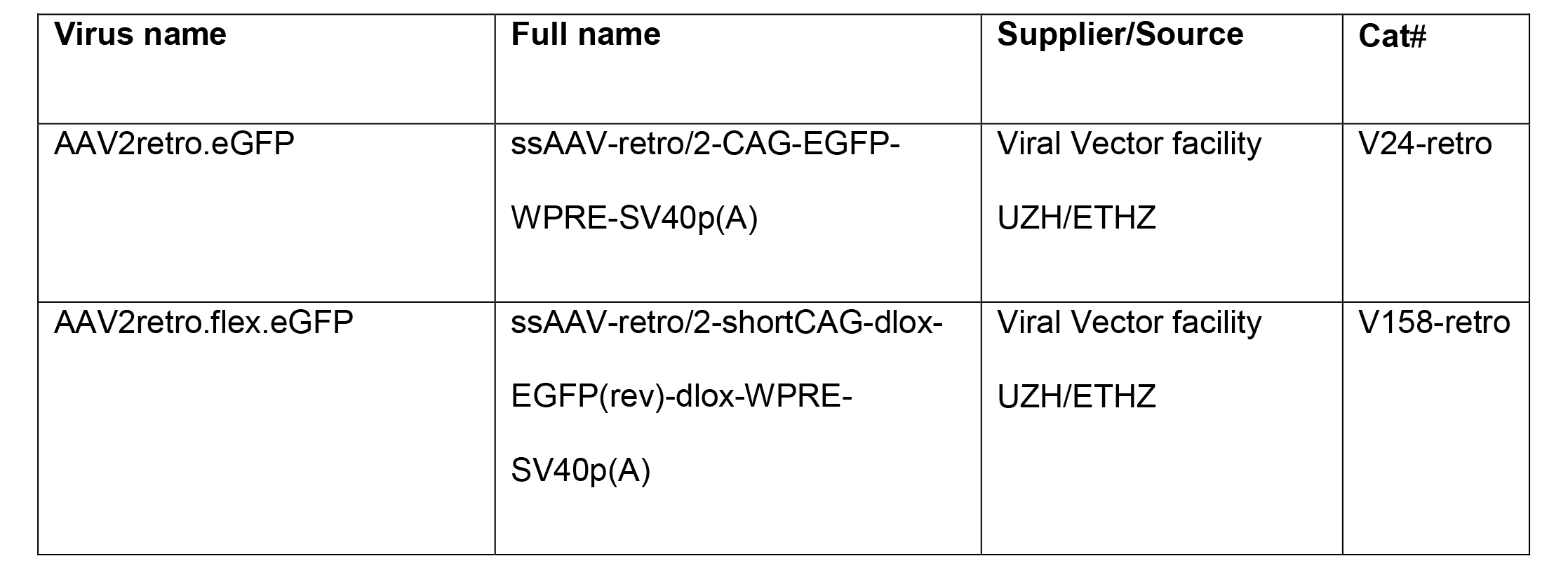

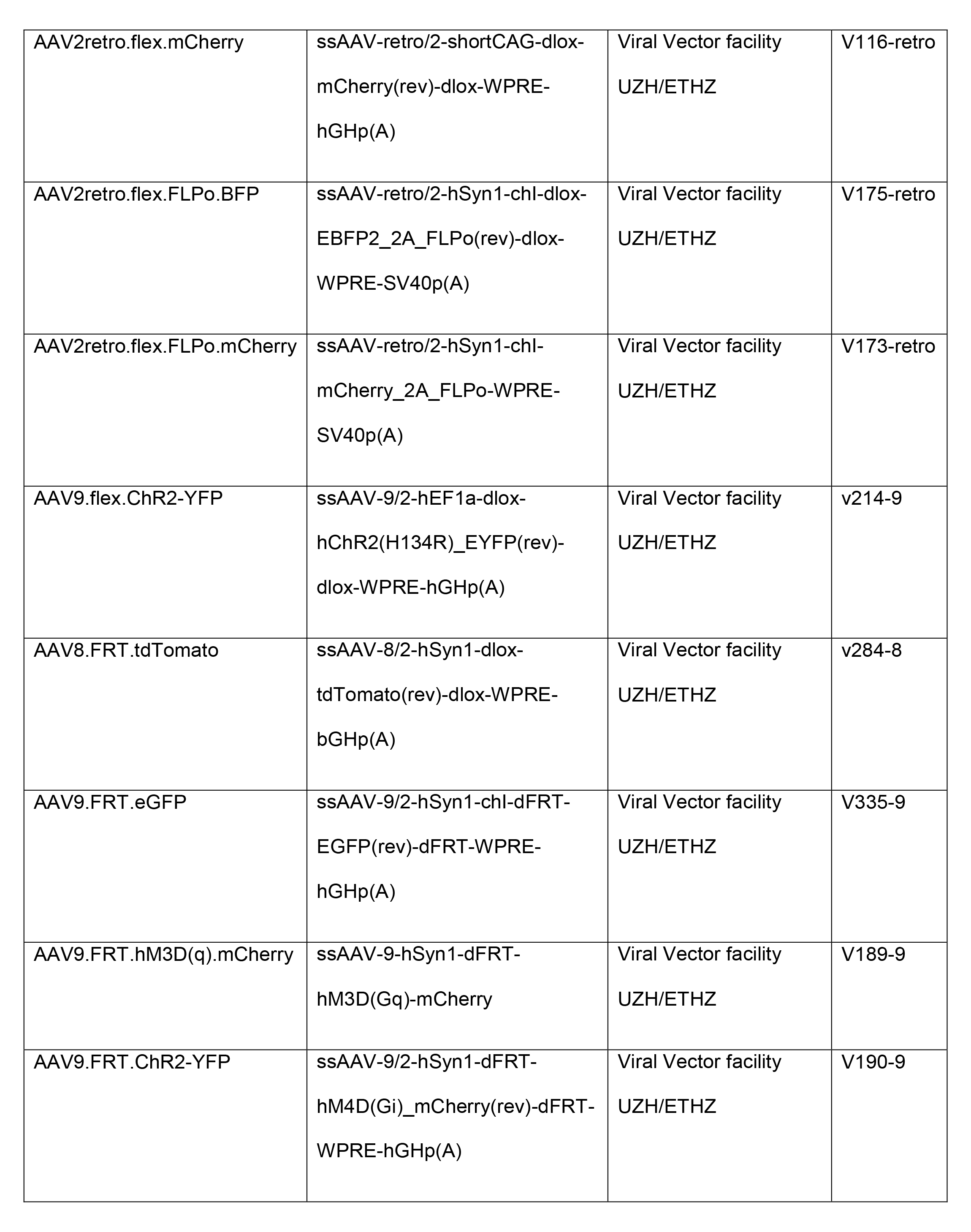

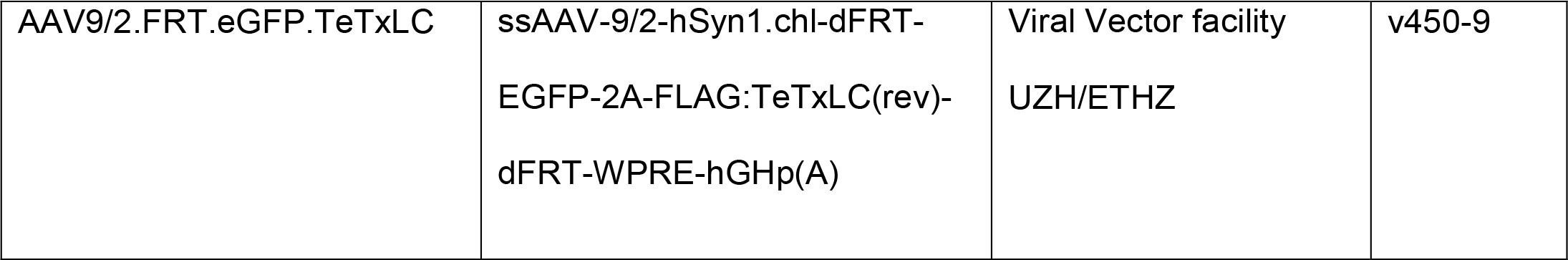

| Reagent | Source | Identifier |
| --- | --- | --- |
| Clozapine-N-oxide | Enzo life sciences | BBL-NS105-0025 |
| Bicuculline-methochloride | Alomone labs | B-137 |
| Strychnine hydrochloride | Sigma Aldrich | S8753-25G |
| 40% acrylamide solution | Thermofisher Scientific | HC2040 |
| Histodenz™ | Sigma Aldrich | D2158 |

| Mouse line | Supplier/source | Reference |
| --- | --- | --- |
| vGAT<sup>cre</sup> | Jackson lab | #016962/ IMSR_JAX:016962 |

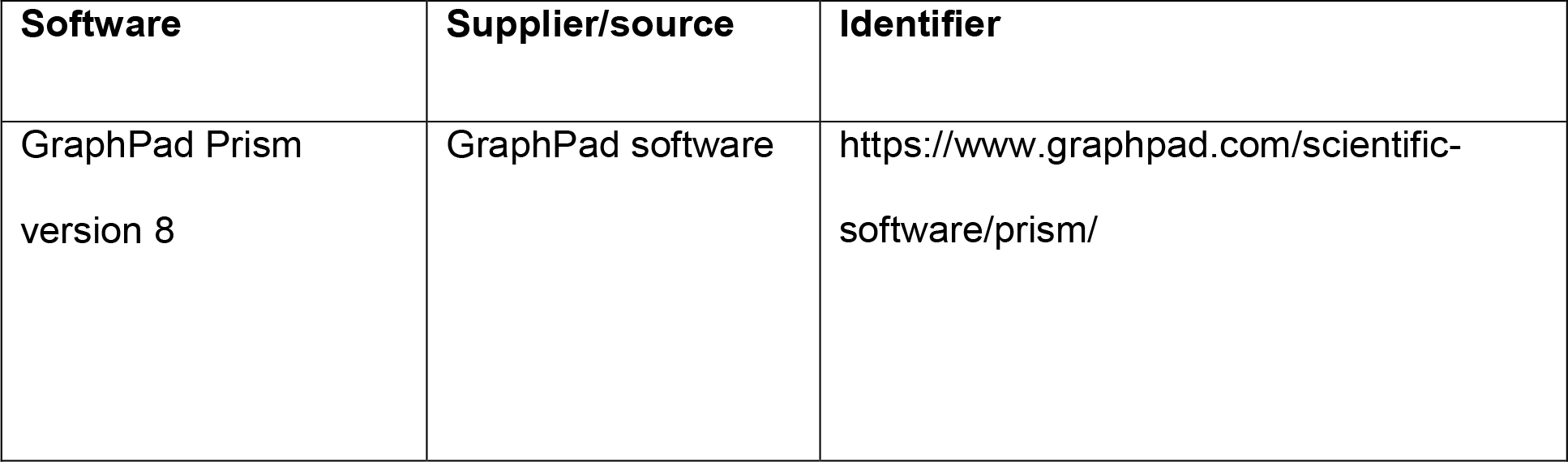

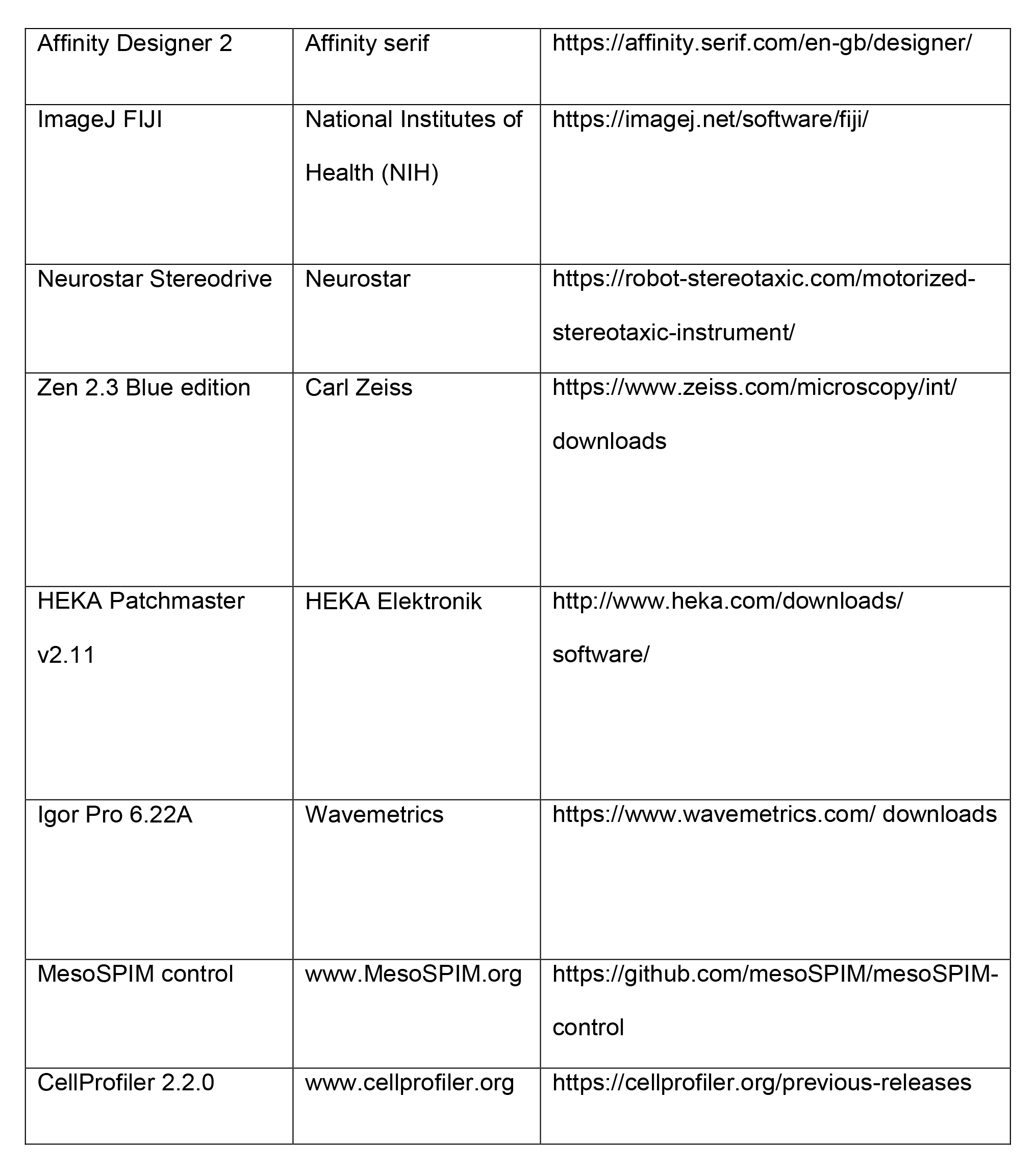
Key resource table.

## Notes

**Conflict of interest: none**

### Competing Interest Statement

The authors have declared no competing interest.

